# Ultrasound Monitoring of Descending Aortic Aneurysms and Dissections in Mice

**DOI:** 10.1101/2020.04.18.048298

**Authors:** Hisashi Sawada, Michael K. Franklin, Jessica J. Moorleghen, Deborah A. Howatt, Masayoshi Kukida, Hong S. Lu, Alan Daugherty

## Abstract

Several modalities, such as computed tomography (CT), magnetic resonance imaging (MRI), and ultrasound, are available to visualize mouse aortas.^1-3^ CT and MRI enable us to obtain reliable images of the aorta and its branches. However, CT requires vascular contrast and MRI is procedurally complex. Thus, these modalities are used only occasionally for in vivo monitoring of mouse studies. High frequency ultrasonography is a common approach for aortic monitoring in mice.^4^ The standard ultrasound approach using a para-sternal view can visualize the aortic root, ascending aorta, and aortic arch, while this approach cannot visualize the descending region due to the presence of lungs and ribs. Therefore, the ability to perform in vivo monitoring of descending aortic diseases in mice has been an impediment. This study reports a para-spinal dorsal approach for ultrasound imaging of mouse descending aortas.

---

Ultrasonography was performed using a Vevo 2100 ultrasound system with a MS550 (40 MHz) transducer in C57BL/6J male mice (9-week-old). Mice were anesthetized by isoflurane as described previously,^4^ and placed on a heated platform (37°C) in a prone position. The transducer was placed on the left edge of the spine at a 45° angle relative to the back (**Figure A**). Aortic images were captured at proximal, mid, and distal regions of the descending thoracic aorta using both long and short axis views (**Figure B**). Aortic imaging of the proximal region was hindered by the interference of the occipital region with the transducer, but descending aortas were visualized clearly in long axis views from the aortic arch to the abdominal aorta. Some cross sectional images in short axis views were not imaged clearly due to artifacts by ribs (**Figure B**). Diameters of the descending aorta were 1.0-1.2 mm (**Figure C**). Subsequently, mice were terminated and aortic diameters were measured in situ to determine concordance with ultrasound measurements (**Figure D**). To maintain aortic patency, optimum cutting temperature (OCT, 300 μl) was injected via the left ventricle using an insulin syringe with a 30G needle after flushing with saline. Aortic diameter was measured at proximal, mid, and distal portions. Bland-Altman plot revealed that ultrasound measurements were consistent with in situ measurements in the proximal region (**Figure D**). However, in the mid and distal regions, ultrasound diameters were larger than diameters in situ. The difference of aortic measurements may be caused by the absence of physiological pressure after euthanasia. Ultrasound is optimal for descending aortic measurements, especially middle to distal regions.

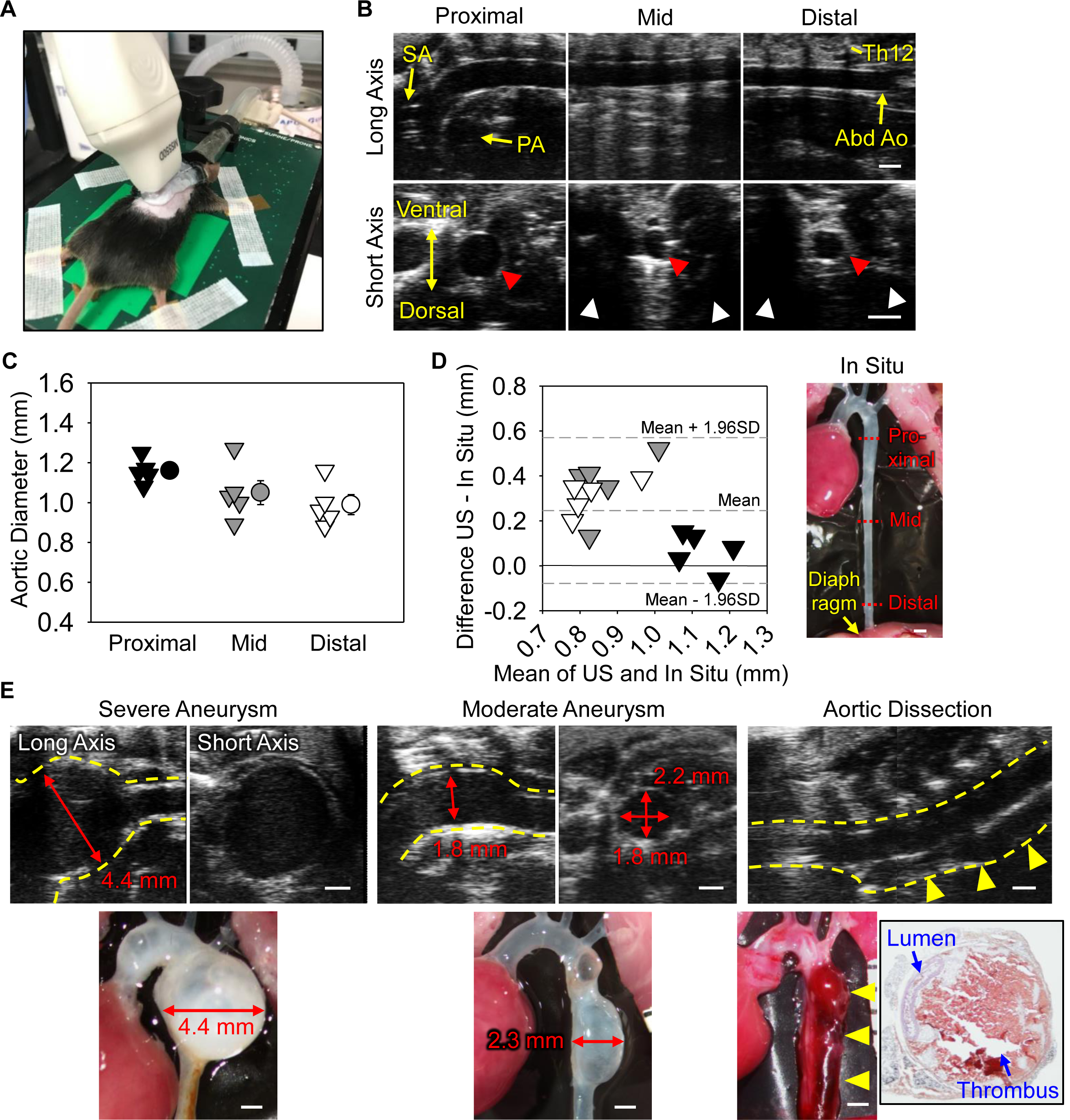
Ultrasound in vivo monitoring of the descending aorta in mice. **(A)** Probe placement for a left paraspinal long axis view. **(B)** Representative long and short axis left para-spinal ultrasound images of non-aneurysmal descending aortas at proximal, mid, and distal regions. SA indicates subclavian artery; PA, pulmonary artery; Abd Ao, abdominal aorta; and Th12, 12th thoracic vertebra. Red and white arrow heads indicate descending aortas and acoustic shadows. **(C)** Aortic diameters at proximal, mid, and distal regions of the descending aorta at mid-systole. n=5. **(D)** Bland-Altman plot shows variation between in situ versus ultrasound measurements at proximal, mid, and distal regions. **(E)** Representative ultrasound images of severe and moderate descending aortic aneurysms and dissection induced by BAPN. Red arrows indicate maximal aortic diameter. Yellow dotted lines and arrow heads indicate the aortic wall and dissected region, respectively. Black box shows hematoxylin-eosin staining of aortic dissection. Scale bar, 1 mm.

We subsequently investigated the application of this approach in mice with aortopathies. To induce pathologies in the descending aorta, β-aminopropionitrile (BAPN, 0.5% wt/vol in drinking water) was administered to male mice for 12 weeks (C57BL/6J, 3-4 weeks old, n=40).^5^ Ultrasonography was performed at 4, 8, and 12 weeks of BAPN administration. It was technically difficult to perform left-para-spinal approach in a mouse with kyphosis. Twenty five mice died during BAPN, and survived mice were terminated when ultrasonography detected luminal dilatation or false lumen. Descending aneurysms were detected ranging from mild to profound that were consistent with in situ measurements (n=5, **Figure E**). Because ultrasonography using the long axis para-spinal view evaluated aortic diameter in a different axis from in situ measurements (ultrasound long axis: anterior-posterior, in situ: left-right), ultrasound aortic diameters should be verified by a short axis view (**Figure E**). Of note, this ultrasound approach also detected aortic dissection (n=2, **Figure E**). A false lumen was observed in the ventral aspect of the descending aorta. Color Doppler did not detect significant blood flow in the false lumen. Aortas were subsequently harvested and stained with hematoxylin and eosin. Hemorrhage was observed at the ventral side in agreement with ultrasound images.

This study demonstrated that ultrasound using the left para-spinal dorsal approach is optimal to monitor progression of aneurysms and dissections in the descending thoracic aorta.

